# Sex-steroid hormones relate to cerebellar structure and functional connectivity across adulthood

**DOI:** 10.1101/2024.06.24.600454

**Authors:** Thamires N. C. Magalhães, Tracey H. Hicks, T. Bryan Jackson, Hannah K. Ballard, Ivan A. Herrejon, Jessica A. Bernard

**Affiliations:** Department of Psychological and Brain Sciences, Texas A&M University, College Station, Texas, United States of America; Vanderbilt Memory & Alzheimer’s Center, Nashville, Tennessee, United States of America; Department of Psychological Sciences, William Marsh Rice University, Houston, Texas, United States of America; Texas A&M Institute for Neuroscience, Texas A&M University, College Station, Texas, United States of America

**Keywords:** aging, sex-specific differences, cerebellar structure, functional connectivity, sex-steroid hormones

## Abstract

Aging involves complex biological changes that affect disease susceptibility and aging trajectories. Although females typically live longer than males, they have a higher susceptibility to diseases like Alzheimer’s, speculated to be influenced by menopause, and reduced ovarian hormone production. Understanding sex-specific differences is crucial for personalized medical interventions and gender equality in health. Our study aims to elucidate sex differences in regional cerebellar structure and connectivity during normal aging by investigating both structural and functional connectivity variations, with a focus on investigating these differences in the context of sex-steroid hormones. The study included 138 participants (mean age = 57(13.3) years, age range = 35-86 years, 54% women). The cohort was divided into three groups: 38 early middle-aged individuals (EMA) (mean age = 41(4.7) years), 48 late middle-aged individuals (LMA) (mean age = 58(4) years), and 42 older adults (OA) (mean age = 72(6.3) years). All participants underwent MRI scans, and saliva samples were collected for sex-steroid hormone quantification (17β-estradiol (E), progesterone (P), and testosterone (T)). We found less connectivity in females between Lobule I-IV and the cuneus, and greater connectivity in females between Crus I, Crus II, and the precuneus with increased age. Higher 17β-estradiol levels were linked to greater connectivity in Crus I and Crus II cerebellar subregions. Analyzing all participants together, testosterone was associated with both higher and lower connectivity in Lobule I-IV and Crus I, respectively, while higher progesterone levels were linked to lower connectivity in females. Structural differences were observed, with EMA males having larger volumes compared to LMA and OA groups, particularly in the right I-IV, right Crus I, right V, and right VI. EMA females showed higher volumes in the right lobules V and VI. These results highlight the significant role of sex hormones in modulating cerebellar connectivity and structure across adulthood, emphasizing the need to consider sex and hormonal status in neuroimaging studies to better understand age-related cognitive decline and neurological disorders.

## 1. Introduction

Aging encompasses a complex biological journey marked by various key features that accrue over time, such as cellular senescence, epigenetic changes, and immune aging, among others [1]. These features, along with their intricate interplay, define the aging process, influencing an individual’s trajectory of aging and susceptibility to diseases [2]. Women typically outlive men, with around 50 men for every 100 women at age 90 [3], due to a complex interplay of biological and sociocultural factors. As the population ages, understanding the factors affecting healthspan, and the length of disease-free life, becomes increasingly important. Despite advancements in health systems, public health campaigns, and health literacy, gender disparities persist. Women show greater susceptibility to diseases like Alzheimer’s [3–5], thought in part to perhaps be related to menopause, which occurs around age 50 and involves a significant reduction in ovarian hormone production [6], including estradiol [5, 7]. Recognizing these sex-specific differences is crucial for achieving gender equity and tailoring personalized medical interventions to address age-related decline and diseases more effectively [2, 8].

In this context, research endeavors have increasingly investigated sex differences, particularly concerning brain structure and resting-state networks across the lifespan [9–11]. An independent component analysis study revealed that young healthy males exhibited stronger connections in the parietal and occipital lobes [9]. At the same time, females showed greater connectivity in the frontal, temporal, and cerebellar regions [9]. These spatially separate brain regions, though distinct, correlate in their activity, forming networks characteristic of the resting state. In other words, the study shows how different brain regions are naturally linked and communicate even when at rest, though the patterns of these interactions differ between the sexes. Goldstone et al. (2016) explored the effects of age and sex on intra- and inter-network FC in cortical networks (default mode (DMN), dorsal attention, and salience). Older adults (OA) showed reduced intra-network FC in the default mode network and increased inter-network FC between the anterior cingulate cortex and dorsal attention network nodes, with these changes being more pronounced in males [10]. Ballard et al. (2022) revealed that network segregation decreased with age, with qualitative differences in network segregation between males and females [11]. Among these investigations, though the cerebellum has been implicated, targeted cerebellar research remains limited.

The cerebellum contributes to both motor and cognitive function and is impacted by both aging and neurodegenerative diseases [12–14]. Stronger connectivity between the cerebellum, the DMN, and striatal pathways is associated with better sensorimotor and cognitive task performance in OA [15–17]. Further, research has reported disrupted FC in elderly populations, particularly in the corticocerebellar and dentate networks [16, 18], as well as in the motor system network [19]. This disrupted FC may underlie some of the motor and cognitive declines observed in aging individuals. Furthermore, distinct resting-state networks have been identified in the dorsal and ventral aspects of the dentate, highlighting the complexity of cerebellar involvement in different functional domains [18].

Another crucial point to consider is the cerebellum’s abundance of estrogen and progesterone receptors [20–22], making it highly sensitive to hormonal fluctuations. During menopause, women experience significant hormonal changes, including a sharp decline in estrogen and progesterone levels [6]. These hormonal changes can affect brain function and structure, particularly in regions with high concentrations of these receptors, such as the cerebellum. Estrogen and progesterone play critical roles in neuroprotection, synaptic plasticity, and cognitive function [23]. The decline of these hormones during menopause may contribute to structural changes in the cerebellum, influencing motor and cognitive functions [14, 24].

Ballard et al. (2022) demonstrated that cerebellar connectivity differences in menopausal females relative to reproductive females are not just related to age, as the patterns differ significantly relative to comparisons in age-matched male groups [11]. That is, reproductive females show greater differences in connectivity between Crus I and Crus II and the medial temporal lobe relative to age matched male groups; however, males show greater differences between Crus I and Crus II and the motor cortex. This finding suggests that the transition to menopause and presumably, the associated hormonal declines are significant for understanding sex differences in aging. However, more research with direct hormonal data is needed to clarify these influences as the work described above was based solely on self-reported menstrual cycles and did not include measures of hormones. As already mentioned, despite the growing body of research, studies examining the effects of sex steroid hormones on cerebellar networks remain limited [25, 26], and conflicting findings persist regarding the potential neuroprotective effects of these hormones on cognitive and brain functions [5, 25, 26]. These hormonal influences suggest that menopause and other periods of hormonal change could significantly affect cerebellar volume, potentially leading to sex-specific aging processes [27].

Although studies consistently show sex differences in cerebellar volume, the patterns of these results vary between different cerebellar lobes, highlighting the need for further research to better understand these complex relationships and the underlying hormonal mechanisms. Raz et al. (2001) [28] and Xu et al. (2000) [29] found greater white matter volume in the posterior cerebellum, as well as greater hemispheric volume and volumetric atrophy in elderly men. Han et al. (2020) [30] found sex differences in multiple hemispheric and vermal cerebellar regions in OA, with significant sex differences over time in left lobule VI and right Crus II longitudinally. Notably, the directionality of these differences varied between lobules. Bernard et al. (2015) observed sex differences in age-cerebellar volume associations, with women (12 to 65 years) showing a more quadratic relationship and men showing a more linear relationship in this age range [31]. The timing of the peak volume in females suggests that perhaps the hormonal changes with menopause may be impacting these age-volume relationships. However, further work is needed to better understand age and sex differences in regional cerebellar volume.

Therefore, the primary objective of our study is to understand sex differences in regional cerebellar structure and connectivity in healthy aging. In addition to investigating structural and functional connectivity differences in males and females, we aim to assess whether these differences are associated with the sex hormones 17ß-estradiol (E), progesterone (P), and testosterone (T). We hypothesize that hormonal fluctuations, especially the decline in E and P during menopause, are associated with specific changes in cerebellar volume and connectivity, potentially accounting for the observed sex differences. By exploring the role of the cerebellum in aging and identifying sex differences in this region, our work seeks to uncover the mechanisms contributing to sex-specific vulnerabilities to age-related decline and disease. Ultimately, our study aims to enhance our understanding of the cerebellum’s role in the aging process and pave the way for future research to develop sex-specific therapeutic strategies that address the unique challenges faced by males and females as they age.

## 2. Methods

### 2.1. Participants

The data used here are part of a larger study on longitudinal changes in the cerebellum across adulthood. One hundred and thirty-eight participants (total n=138, 57±13.3 years, range=35-86 years at baseline, 54% females) were included in this study, comprising 38 individuals in early middle age (EMA) (41±4.7 years), 48 in late middle age (LMA) (58±4 years), and 42 older adults (OA) (72±6.3 years). All participants underwent a series of cognitive and motor tasks and provided saliva samples for hormone quantification during the assessment (details on sample collection are provided below). Subsequently, they returned for a magnetic resonance imaging (MRI) session approximately two weeks later. However, due to unforeseen delays related to the COVID-19 pandemic, the time between the two sessions varied among participants (mean 39.0 days ± 21.4 days). For the analyses presented here, we focused solely on hormonal and brain imaging data. However, our group has previously conducted a study addressing both brain imaging data and behavior [32].

Exclusion criteria for participation in the study included a history of neurological disease, stroke, or formal diagnosis of psychiatric illness (e.g., depression or anxiety), contraindications for the brain imaging environment, and the use of hormone therapy (HTh) or hormonal contraceptives (including intrauterine devices (IUDs), possible use of continuous birth control (oral), and no history of hysterectomy in the past 10 years). These exclusions were implemented to assess the effects of normative endocrine aging on healthy adult females. All study procedures were approved by the Institutional Review Board at Texas A&M University, and written informed consent was obtained from each participant before any data collection commenced.

### 2.2. Hormone quantification

We followed the methodology described in our recent work [14] for hormonal analyses, and to ensure clarity and replicability, we have shared those methods directly here. Before collecting the saliva sample, participants were asked to refrain from consuming alcohol 24 hours prior and eating or drinking 3 hours before their first study session to avoid exogenous influences on hormone levels. Participants were also screened for oral disease or injury, use of substances such as nicotine or caffeine, and prescription medications that may impact the saliva pH and compromise samples. Participants were asked to rinse their mouths with water for 10 minutes before providing a saliva sample to clear out any residue.

Samples were then collected in pre-labeled cryovials provided by Salimetrics (https://salimetrics.com/saliva-collection-training-videos/) using the passive drool technique. For our study, participants were asked to supply 1mL of saliva, after which samples were immediately stored in a −80° Celsius bio-freezer for stabilization. Assays were completed by Salimetrics to quantify 17β-estradiol, progesterone, and testosterone levels for each participant. The amount of saliva collected was sufficient to detect 17β-estradiol at a high sensitivity threshold of 0.1 pg/mL[33], along with 5.0 pg/mL and 1.0 pg/mL thresholds for progesterone and testosterone, respectively.

### 2.3. Imaging acquisition

Participants underwent structural and resting-state MRI using a Siemens Magnetom Verio 0 Tesla scanner with a 32-channel head coil. Structural MRI included a high-resolution T1-weighted 3D magnetization prepared rapid gradient multi-echo (MPRAGE) scan (TR = 2400 ms; acquisition time = 7 minutes; voxel size = 0.8 mm^3^) and a high-resolution T2-weighted scan (TR = 3200 ms; acquisition time = 5.5 minutes; voxel size = 0.8 mm^3^), both with a multiband acceleration factor of 2. Resting-state imaging comprised four blood-oxygen-level-dependent (BOLD) functional connectivity (fMRI) scans with a multiband factor of 8, 488 volumes, TR of 720 ms, and 2.5 mm^3^ voxels. Each fMRI scan lasted 6 minutes, totaling 24 minutes of resting-state imaging, with scans acquired in alternating phase encoding directions (i.e., two anterior-to-posterior scans and two posterior-to-anterior scans). Participants were instructed to lie still with their eyes open, fixating on a central cross during fMRI scans. The total image acquisition time was approximately 45 minutes, including a 1.5-minute localizer scan. Scanning protocols were adapted from the Human Connectome Project [34] and the Center for Magnetic Resonance Research at the University of Minnesota to ensure future data sharing and reproducibility.

#### 2.3.1. Functional imaging processing and connectivity analysis

The images were converted from DICOM to NIFTI format and organized according to the Brain Imaging Data Structure (BIDS, version 1.6.0) using the bidskit docker container (version 2021.6.14, https://github.com/jmtyszka/bidskit). A single volume was extracted from two oppositely coded BOLD images to estimate B0 field maps using the split tool from the FMRIB Software Library (FSL) package [35]. Subsequently, anatomical, and functional images were preprocessed using fMRIPrep (version 20.2.3, https://fmriprep.org/), which involves a series of automated procedures to prepare fMRI data for further analysis. This includes aligning the functional volume with the anatomical image for accurate spatial registration, correcting motion to account for participant movement during image acquisition, correcting fieldmap distortions caused by the non-uniform magnetic field, segmenting the anatomical image into distinct tissues (e.g., gray matter, white matter, cerebrospinal fluid (CSF)), stripping the skull from the anatomical image to improve segmentation quality and reduce artifacts, normalizing the data to a common space for cross-participant or cross-study comparisons, aligning motion-corrected functional volumes with the normalized anatomical image for functional-anatomical integration, and applying spatial smoothing to enhance the signal-to-noise ratio and reduce artifacts. We continued the rest of the analyses using the CONN toolbox (version 21a) [36]. A 0.008-0.099 Hz band-pass filter is applied to remove high-frequency noise. The threshold for global signal z-values was set to 3, and the motion correction threshold was set to 0.5 mm. Motion data along 6 axes and frame outliers, eliminated during denoising to adhere to the global mean, were included as first-level covariates.

Regions of interest (ROIs) – lobule masks – included Crus I, Crus II, lobule V, lobules I-IV, and vermis VI – right hemisphere, with masks created using the SUIT atlas [37], as described in previous studies [16, 38, 39]. These ROIs are part of broader neural networks connecting the cerebellum and cerebral cortex[38]. Exploring their connectivity can enhance our understanding of the cerebellum’s role in both motor and non-motor functions. For instance, Crus I and Crus II are linked with higher-order cognitive processes and motor planning, while lobule V and lobules I-IV are crucial for integrating sensory input with motor commands, essential for coordinated movement and proprioception [14, 40]. The vermis, particularly vermis VI, is involved in emotional regulation and connects to the limbic system [41]. Focusing on the right hemisphere allows for better observation of functions related to attention and visuospatial processing [42]. Group-level whole-brain seed-to-voxel analyses were performed to examine connectivity patterns in relation to age. We employed standard settings for cluster-based inferences using parametric statistics based on random field theory. We used an initial voxel threshold at p<.001 along with a cluster threshold set at p < .05, with a false discovery rate (FDR) correction.

#### 2.3.2. Structural image processing

As described earlier, we began by converting high-resolution T1-weighted 3D images from DICOM to NIFTI format. Subsequently, we used FastSurfer software [43, 44] (https://github.com/Deep-MI/FastSurfer) in conjunction with the latest version of FreeSurfer (version 7.4.1) (https://github.com/freesurfer/freesurfer) for our volumetric segmentation analysis. FastSurfer, an open-source tool, is specifically designed to automatically segment cortical gray matter and calculate key morphometric measurements including cortical thickness, surface area, and volume. To enhance our structural analysis, we integrated CerebNet [45], a recent addition to the FastSurfer and FreeSurfer packages. CerebNet focuses on the analysis of the cerebellum and its subregions, providing tools for cerebellar segmentation, surface reconstruction, and morphometric measurements. CerebNet is a fully automated, extensively validated deep-learning method for cerebellar lobular segmentation. It combines FastSurferCNN, a UNet-based 2.5D segmentation network, with extensive data augmentation techniques such as realistic non-linear deformations. This approach increases the anatomical variety and eliminates the need for additional preprocessing steps like spatial normalization or bias field correction. CerebNet has demonstrated high accuracy, surpassing state-of-the-art approaches in cerebellar segmentation [45].

The image analysis workflow includes pre-processing, removal of non-brain tissues, segmentation of white matter, gray matter, and CSF, and generation of a three-dimensional mesh representing the cortical surface. A post-processing step refines the surface mesh, corrects artifacts, divides the surface into specific anatomical regions, and calculates morphometric measurements for each region of interest (Crus I, Crus II, lobule V, lobule I-IV, and lobule VI). The output includes files containing calculated morphometric measurements and total intracranial volume (TIV), and these results were exported to an Excel file for further statistical analysis.

### 2.4. Structural data statistical analysis

For the statistical analysis of structural data, we used IBM SPSS software (version 29, SPSS Inc., Chicago, IL, USA). The statistical analysis was conducted using the General Linear Model (GLM) to examine the effects of age group (EMA, LMA, and OA) and sex (female and male) on the measurements of five cerebellar subregions (Right I-IV, Right V, Right VI, Right Crus I, and Right Crus II). For each cerebellar subregion, the GLM was specified with the dependent variable being the measurement of the subregion (e.g., volume (mm^3^)). The independent variables included age group, sex, and the interaction between age group and sex, we also included TIV as a covariate in this model. Tests of between-subjects effects were conducted to examine the main effects and interaction effects. Post-hoc Bonferroni comparisons were conducted to further investigate significant interactions, with significance set at p < 0.05. The results were interpreted by examining the estimated marginal means and the significance of the main and interaction effects.

A similar GLM model was performed to examine the effects of age group and sex on hormone levels (pg/mL) (E, P, and T). Graphs were generated to visualize the interaction effects using the estimated marginal means, providing a clear representation of how the measurements of the cerebellar subregions and hormone levels varied across different age groups and sexes.

After conducting group comparisons, we employed linear regression models to examine whether hormone levels predict structural changes in specific cerebellar subregions. For these analyses, ROI volumes were normalized to each subject’s total intracranial volume (TIV), resulting in a percentage to account for individual differences in brain size. Each model included the hormone level as the independent variable and the TIV-normalized volume of the ROI as the dependent variable.

## 3. Results

### 3.1. Functional connectivity

When examining interactions between age and connectivity in males and females we found several relationships. (Figure 1, Table 1). Specifically, there was significantly higher connectivity between Lobules I-IV and the cuneus in males as compared to females with increased age (Figure 1A; Table 1). Further, lower connectivity was demonstrated between both Crus I and Crus II with the precuneus in males as compared to females with increased age (Figure 1C; Table 1). However, higher connectivity was shown between Lobule V and the calcarine cortex in females as compared to males with greater age (Figure 1B; Table 1). Lastly, lower connectivity between Vermis VI and Crus I was demonstrated in males as compared to females with increased age (Figure 1D; Table 1).

**Figure 1.**
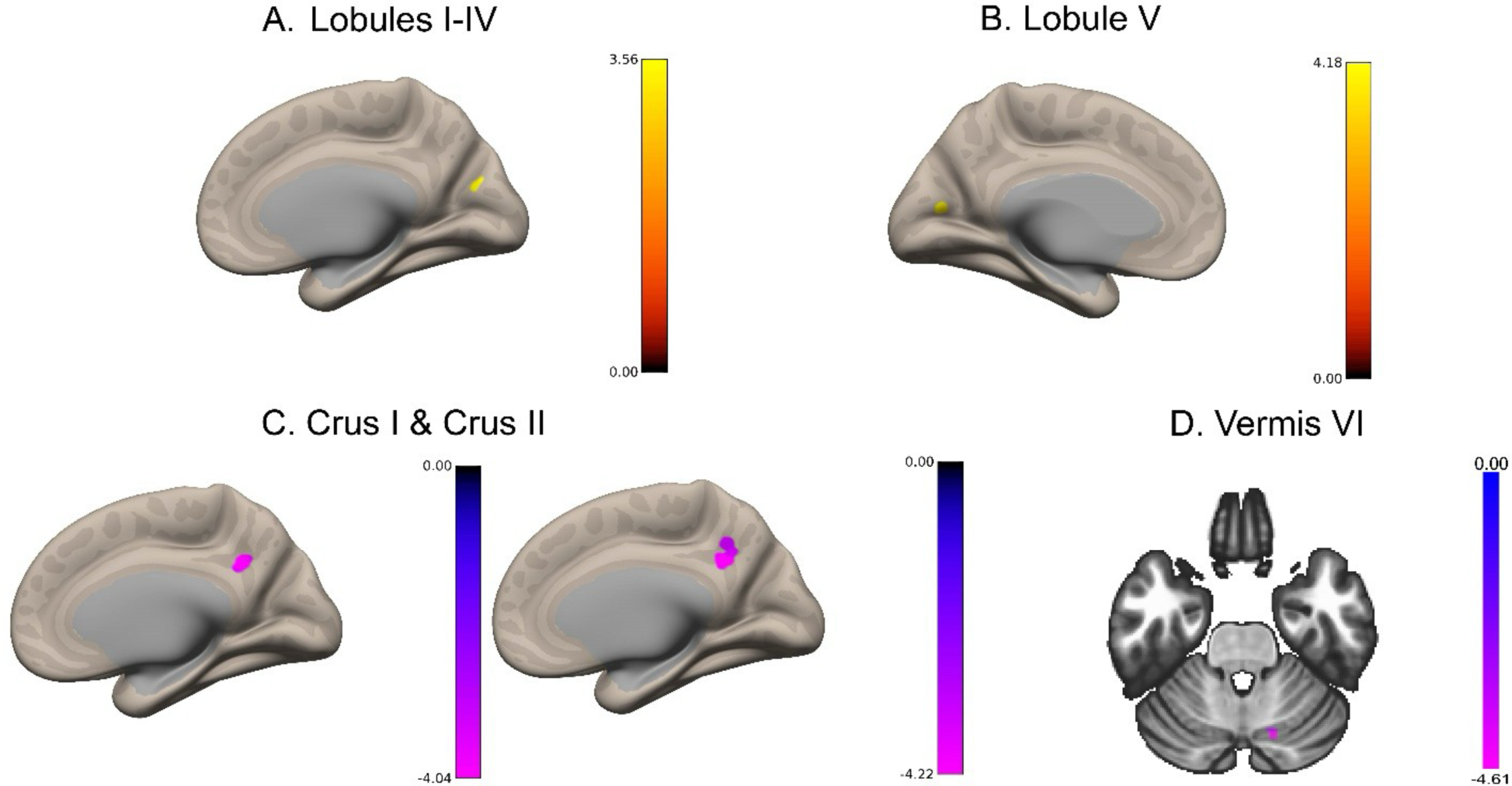
Bidirectional cortical functional connectivity (FC) relationships in males as compared to females (M>F) with increased age across ROIs. **A.** yellow displays greater FC between lobules I-IV and the cuneus in males as compared to females with increased age. **B.** yellow represents greater FC between lobule V and the calcarine cortex in males as compared to females with increased age. **C.** purple displays lower FC between Crus I (left) and Crus II (right) and the precuneus in males as compared to females with increased age. **D.** purple represents lower FC between vermis VI and Crus I in males as compared to females with increased age.

**Table 1.** Coordinates of cerebellar regions showing significant correlations in cortical FC with age by sex interaction.

| Cortical Functional Connectivity Correlations<br>with Age by Sex Interaction<br>Male>Female |  |  |  |  |  |  |
| --- | --- | --- | --- | --- | --- | --- |
| ROI | X | Y | Z | Cluster<br>Size | T(134) | pFDR |
| **Lobules I-IV** |  |  |  |  |  |  |
| Cuneus | 0 | -72 | 26 | 18 | 5.10 | 0.0198 |
| **Crus I** |  |  |  |  |  |  |
| Precuneus | 16 | -56 | 36 | 22 | -4.97 | 0.0081 |
| **Crus II** |  |  |  |  |  |  |
| Precuneus | 12 | -52 | 32 | 39 | -5.62 | 0.0001 |
| **Lobule V** |  |  |  |  |  |  |
| Calcarine cortex | -2 | -74 | 12 | 18 | 4.66 | 0.0296 |
| **Vermis VI** |  |  |  |  |  |  |
| Crus I | 16 | -72 | -28 | 17 | -5.30 | 0.0408 |

### 3.2. Hormone Level Associations Across Participants

Functional connectivity relationships with 17β-estradiol were revealed in Crus I and Crus II across all participants (Figure 2; Table 2). Specifically, there was significantly higher connectivity between Crus I and the superior parietal gyrus with increased 17β-estradiol levels (Figure 2A; Table 2). Higher connectivity was also demonstrated between Crus II and the cuneus with greater 17β-estradiol levels (Figure 2B; Table 2). There were no significant associations between Lobules I-IV, Lobule V, or Vermis VI (pFDR > .05).

**Figure 2.**
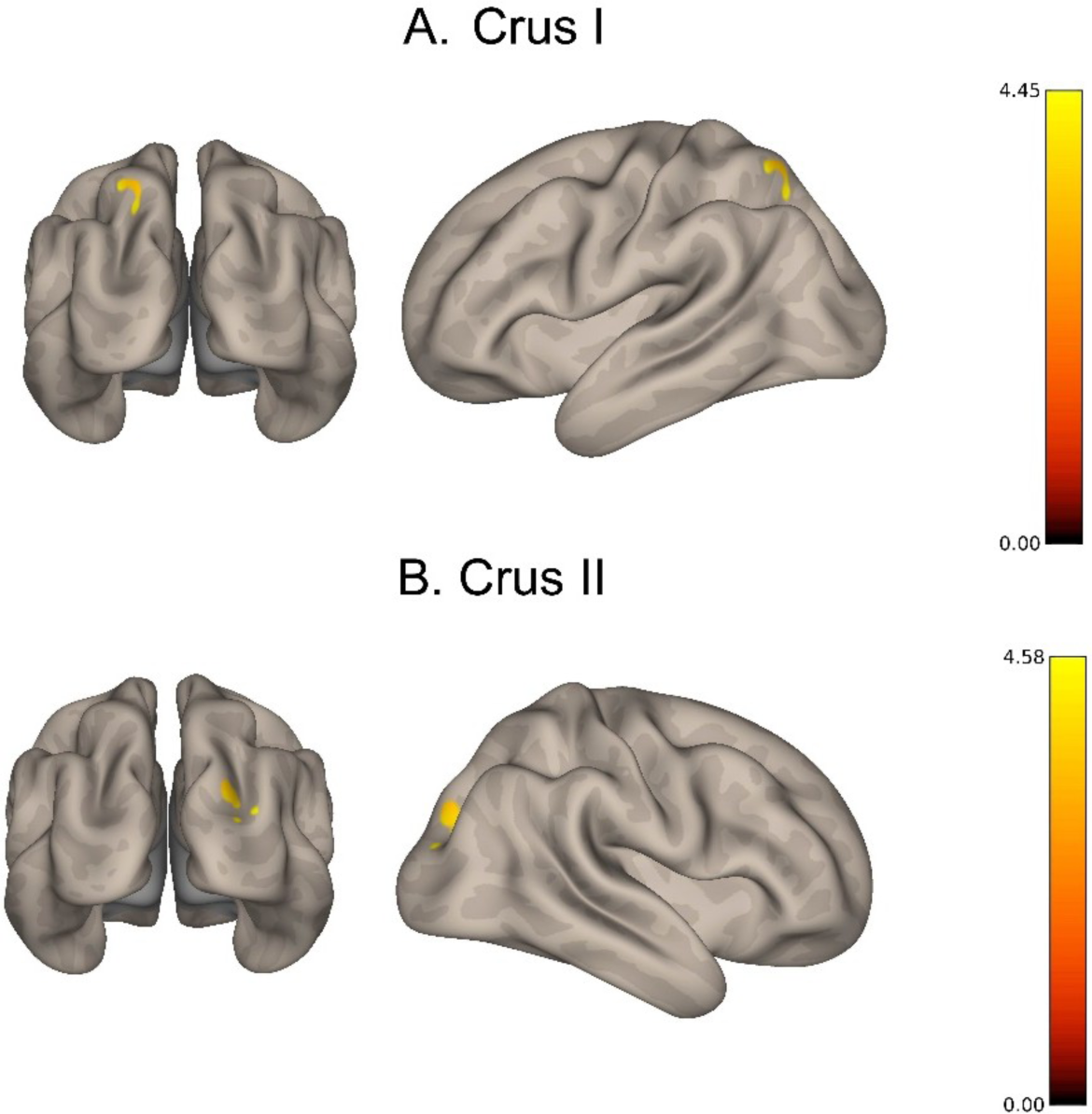
Higher cortical connectivity associations in Crus I and Crus II were observed with higher levels of 17β-estradiol across participants. **A.** yellow displays greater FC between Crus I and the superior parietal gyrus with increased 17β-estradiol. **B.** yellow represents greater FC between Crus II and the cuneus with increased 17β-estradiol.

**Table 2.** Coordinates of cerebellar regions showing significant correlations in cortical FC with 17β-estradiol.

| Cortical Functional Connectivity Correlations<br>with Estradiol Across Participants by ROI |  |  |  |  |  |  |
| --- | --- | --- | --- | --- | --- | --- |
| ROI | X | Y | Z | Cluster Size | T(127) | pFDR |
| **Crus I** |  |  |  |  |  |  |
| Superior Parietal gyrus | -22 | -58 | 52 | 22 | 5.44 | 0.0052 |
| **Crus II** |  |  |  |  |  |  |
| Cuneus | 30 | -82 | 20 | 23 | 5.60 | 0.0033 |

Investigations of testosterone and connectivity across all participants revealed several associations. There was significantly higher connectivity between Lobules I-IV and both the angular gyrus and the lingual gyrus (Figure 3A; Table 3) with increased testosterone levels across participants. Lower connectivity was demonstrated between Crus I and both the middle temporal gyrus and the superior frontal gyrus with greater testosterone levels across all participants (Figure 3B; Table 3). In Lobule V there was lower connectivity with the the inferior frontal gyrus with greater testosterone levels across participants (Figure 3C; Table 3) and similar patterns were seen with Vermis VI and the middle temporal gyrus with greater testosterone levels (Figure 3D; Table 3). There were no significant associations between Crus II and testosterone (pFDR > .05).

**Figure 3.**
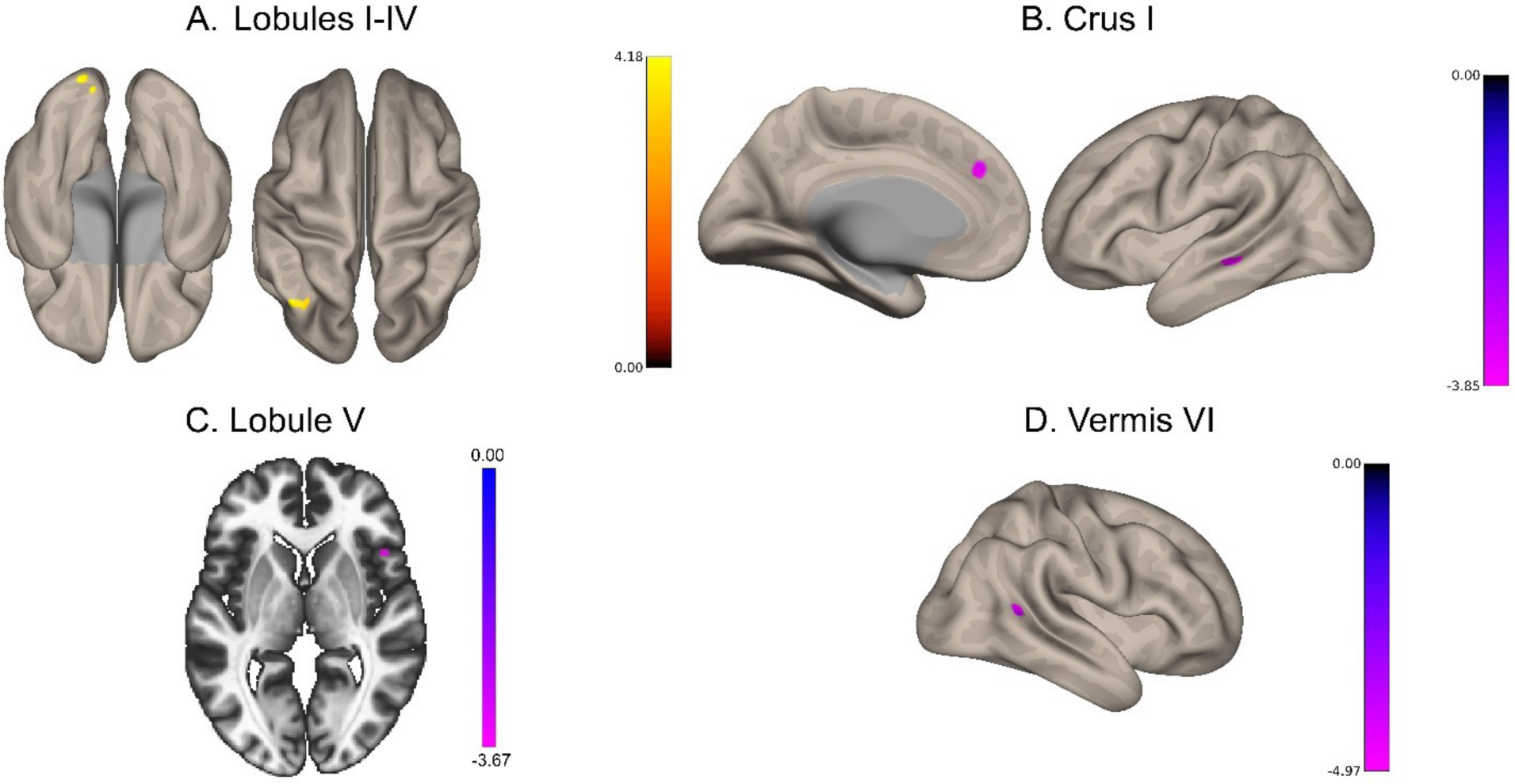
Bidirectional cortical FC associations in ROIs with greater levels of testosterone. **A.** yellow displays greater FC between lobules I-IV and both the angular and lingual gyri with increased testosterone. **B.** purple represents lower FC between Crus I and both the superior frontal and middle temporal gyri with increased testosterone. **C.** purple displays lower FC between lobule V and the inferior frontal gyrus with increased testosterone. **D.** purple represents lower FC between vermis VI and the middle temporal gyrus with increased testosterone.

**Table 3.** Coordinates of cerebellar regions showing significant correlations between cortical FC and testosterone.

| Cortical Functional Connectivity Correlations<br>with Testosterone Across Participants by ROI |  |  |  |  |  |  |
| --- | --- | --- | --- | --- | --- | --- |
| ROI | X | Y | Z | Cluster<br>Size | T(132) | pFDR |
| **Lobules I-IV** |  |  |  |  |  |  |
| Angular gyrus | -38 | -68 | 50 | 17 | 5.50 | 0.0206 |
| Lingual gyrus | -14 | -96 | -18 | 14 | 5.15 | 0.0282 |
| **Crus I** |  |  |  |  |  |  |
| Middle Temporal gyrus | -54 | -32 | -8 | 20 | -5.40 | 0.0199 |
| Superior Frontal gyrus,<br>medial part | 0 | 42 | 32 | 15 | -5.38 | 0.0459 |
| **Lobule V** |  |  |  |  |  |  |
| Inferior Frontal gyrus,<br>opercular part | 48 | 18 | 2 | 18 | -4.64 | 0.0249 |
| **Vermis VI** |  |  |  |  |  |  |
| Middle Temporal gyrus | 56 | -52 | 12 | 20 | -5.13 | 0.0186 |

Analyses were conducted for male and female participants separately to better parse the observed associations between lobular connectivity and hormone levels.

Male participants showed significant positive associations between lobule V and the middle occipital gyrus with increased progesterone levels (Figure 4A; Table 4). Males also displayed greater connectivity between lobules I-IV and the middle temporal gyrus with increased progesterone levels (Figure 4B; Table 4). Further, higher connectivity was shown between Crus I and the inferior frontal gyrus with increased progesterone levels (not visualized); however, lower connectivity was shown between Crus I and the middle temporal, middle frontal, and inferior frontal gyri, as well as the putamen and anterior cingulum with higher progesterone levels (Figure 4C; Table 4). There were no significant associations between Crus II and Vermis VI and progesterone levels with connectivity (pFDR > .05). There were no associations between testosterone or 17β-estradiol levels and ROI connectivity in males (pFDR > 0.05).

**Figure 4.**
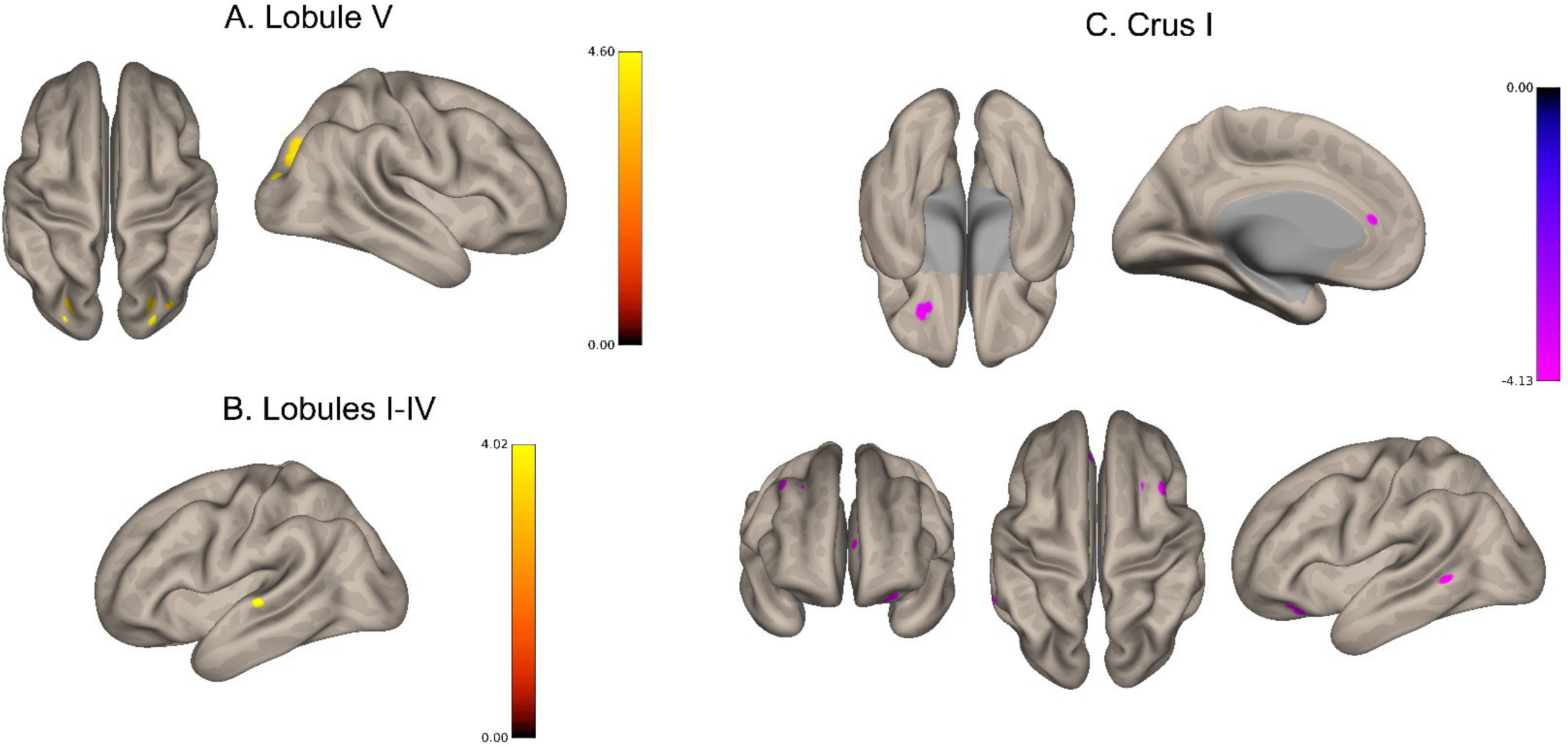
Bidirectional cortical FC relationships associations in ROIs with higher levels of progesterone in males. **A.** yellow displays greater FC between lobule V and the middle occipital gyrus with increased progesterone. **B.** yellow represents greater FC between lobules I-IV and the middle temporal gyrus with increased progesterone. **C.** purple displays lower FC between Crus I and both the middle and inferior frontal gyri, the putamen, the anterior cingulum, and the middle temporal gyrus with increased progesterone.

**Table 4.** Coordinates of cerebellar regions showing significant correlations in cortical FC with progesterone in male participants.

| Cortical Functional Connectivity Correlations<br>with Progesterone in Males by ROI |  |  |  |  |  |  |
| --- | --- | --- | --- | --- | --- | --- |
| ROI | X | Y | Z | Cluster<br>Size | T(60) | pFDR |
| **Lobules I-IV** |  |  |  |  |  |  |
| Middle Temporal gyrus | -64 | -22 | 2 | 19 | 5.48 | 0.0103 |
| **Crus I** |  |  |  |  |  |  |
| Middle Frontal gyrus | 46 | 16 | 52 | 35 | -6.56 | 0.0003 |
| Putamen | 26 | -6 | 6 | 27 | -6.02 | 0.0013 |
| Inferior frontal gyrus,<br>orbital part | -22 | 38 | 0 | 14 | 5.78 | 0.0468 |
| Middle Frontal gyrus | 28 | 16 | 40 | 13 | -4.50 | 0.0468 |
| Anterior Cingulum | -4 | 36 | 12 | 12 | -4.23 | 0.0468 |
| Middle Temporal gyrus | -64 | -52 | 2 | 12 | -4.42 | 0.0468 |
| Inferior Frontal gyrus,<br>orbital part | -24 | 34 | -12 | 12 | -5.73 | 0.0468 |
| **Lobule V** |  |  |  |  |  |  |
| Middle Occipital gyrus | 30 | -84 | 22 | 46 | 6.42 | <0.0001 |
| Middle Occipital gyrus | -24 | -84 | 20 | 28 | 5.49 | <0.0001 |
| Middle Occipital gyrus | 30 | -78 | 32 | 16 | 5.95 | 0.0154 |

When investigating females separately, the associations with hormones were limited to testosterone only. Female participants demonstrated significantly higher connectivity between Crus I and the inferior temporal gyrus with higher testosterone levels (not visualized); however, lower connectivity was seen between Crus I and the parahippocampal gyrus with higher testosterone levels (Figure 5A; Table 5). Stronger connectivity was also shown between Crus II and the inferior temporal gyrus with higher testosterone levels (Figure 5B). Vermis VI connectivity with Lobule VII was positively associated with testosterone levels; but, lower connectivity was shown between Vermis VI and the superior temporal gyrus in females (Figure 5C; Table 5) with higher testosterone levels. There were no significant associations between Lobules I-IV and Lobule V and testosterone levels with connectivity in females (pFDR > .05).

**Figure 5.**
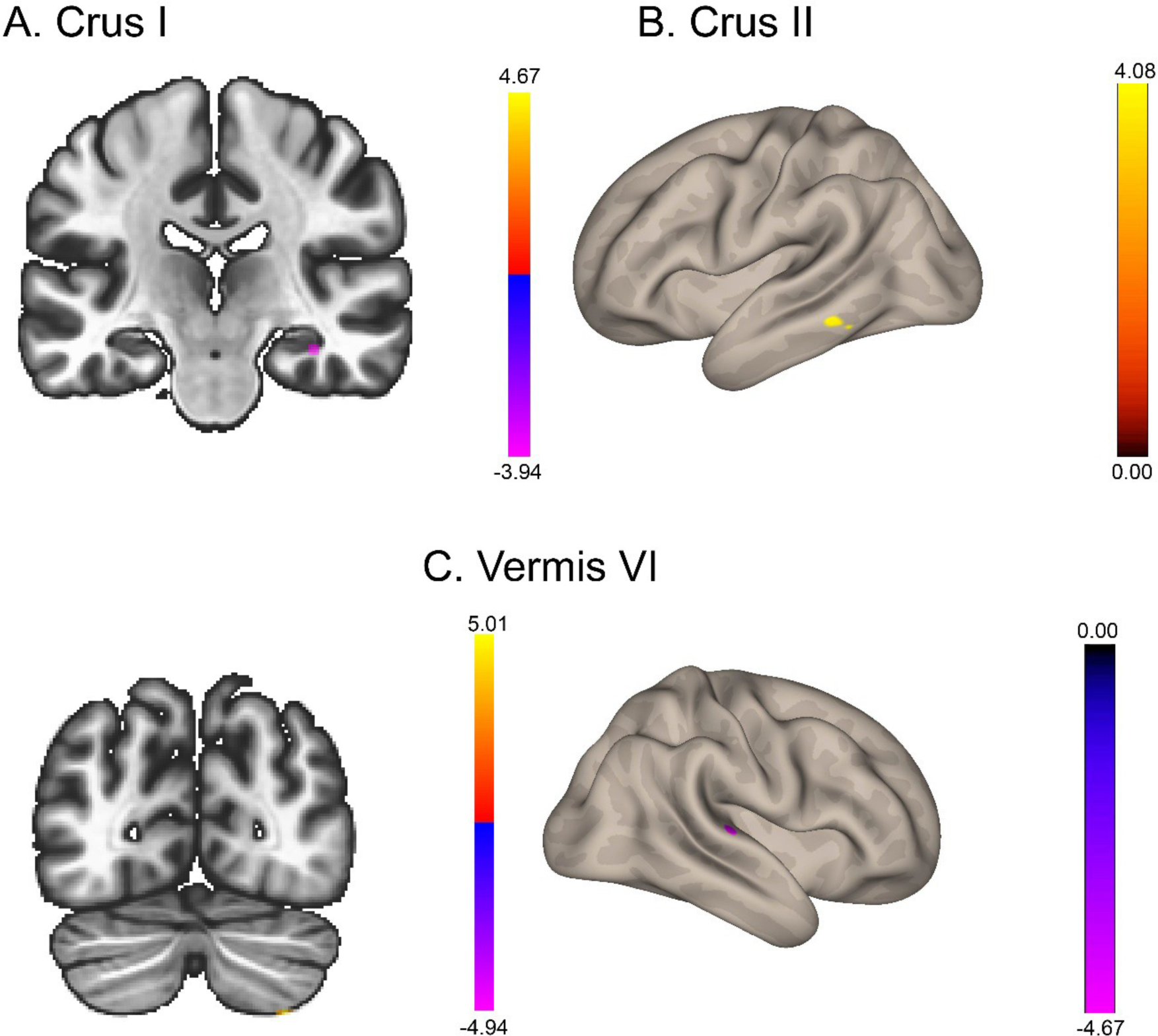
Bidirectional cortical FC relationships between ROIs with higher levels of testosterone in females. **A.** purple displays lower FC between Crus I and the parahippocampal gyrus with increased testosterone. **B.** yellow represents greater FC between Crus II and the inferior temporal gyrus with increased testosterone. **C.** yellow shows greater FC between vermis VI and lobule VII with increased testosterone (left); whereas, purple displays lower FC between vermis VI and superior temporal gyrus with increased testosterone (right).

**Table 5.**
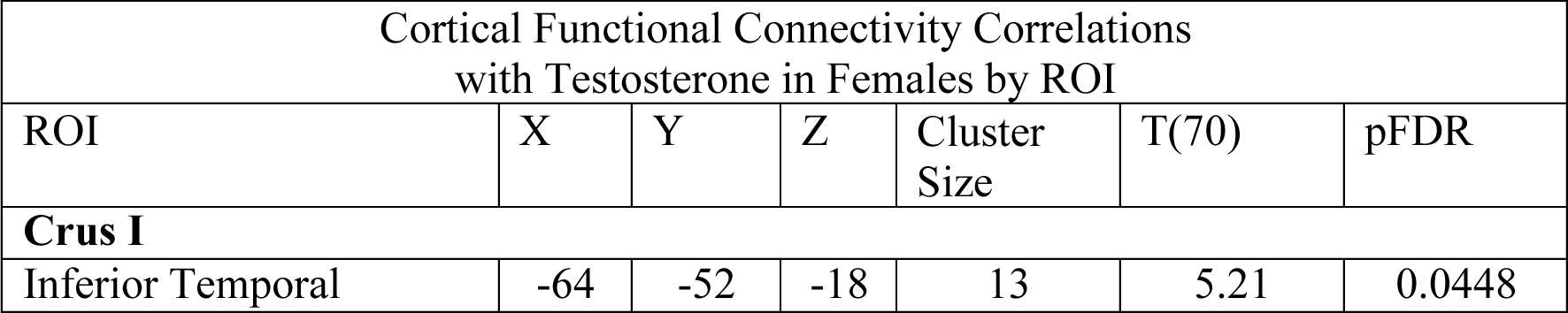
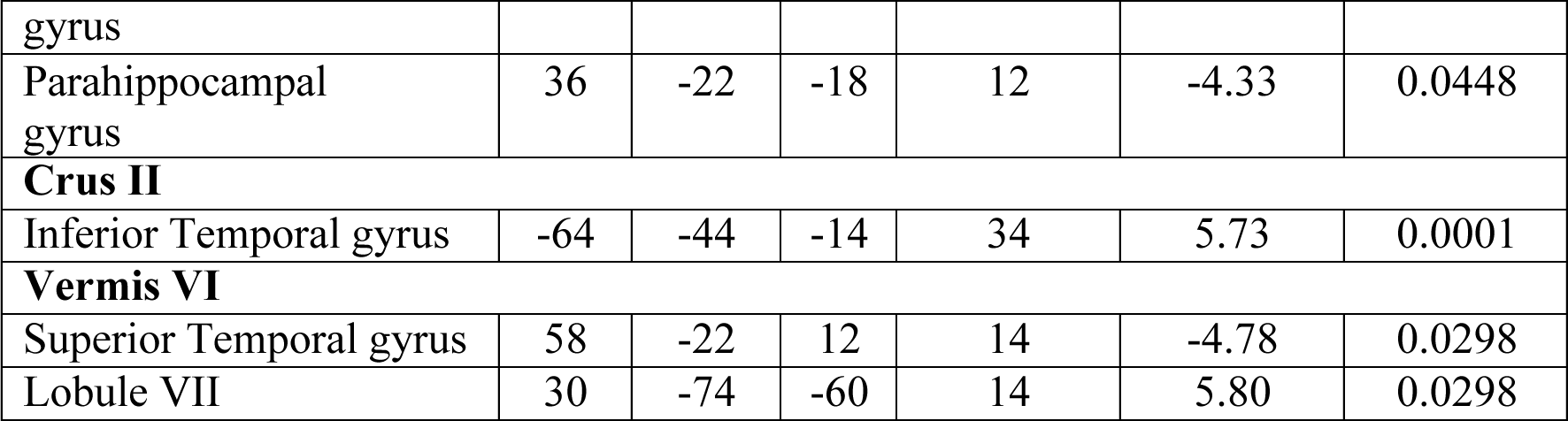
Coordinates of cerebellar regions showing significant correlations in cortical FC with testosterone in female participants.

### 3.3. Structural analysis results

In our general model, we found that sex and age group significantly impact cerebellar subregions, although the interaction between sex and age was not significant, as shown in **Table 6**. Considering the significant results regarding sex, we conducted pairwise comparisons using the Bonferroni post hoc to compare males and females, we observed that males have larger volumes than females in the right cerebellar subregions I-IV and V, as represented in **Figure 7**. For more statistical details, see **Table 7**.

**Figure 7.**
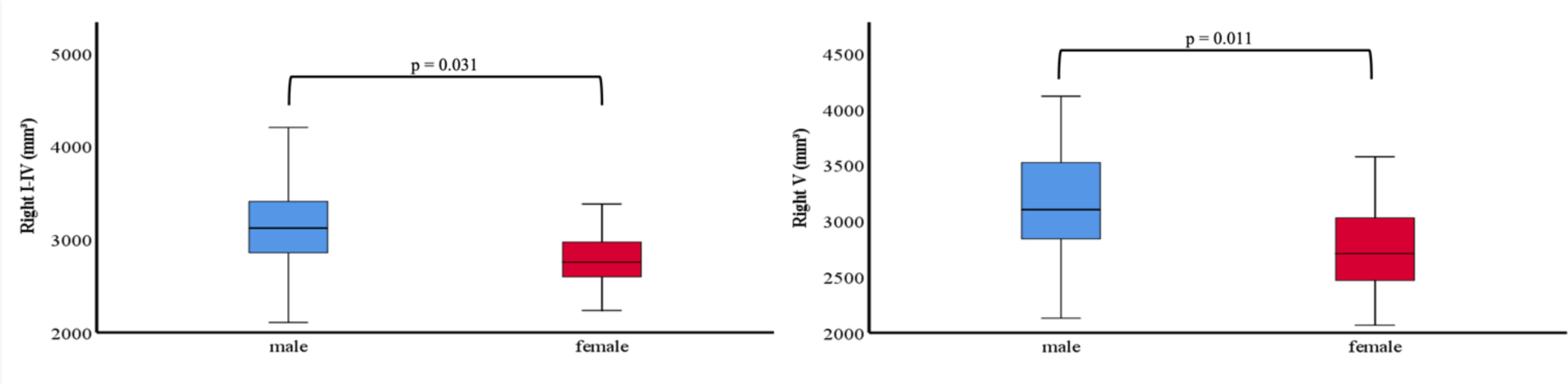
There are significant differences in cerebellar subregion volumes between males and females. For males, the standard error (SE) is 4.15 for the right I-IV and 4.46 for the right V, while for women, the SE is 3.88 for the right I-IV and 4.16 for the right V.

**Table 6.**
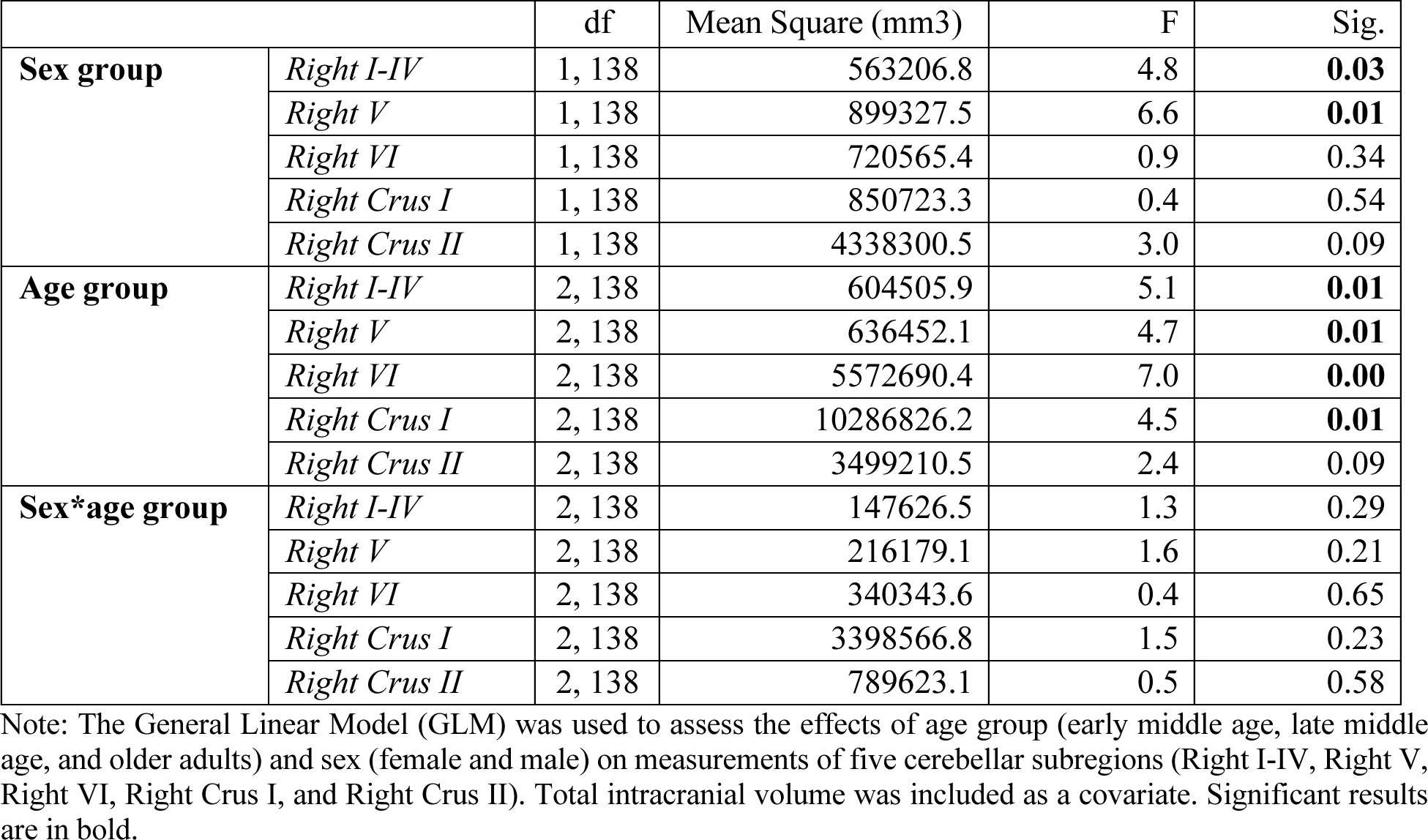
Tests of between-subjects cerebellar subregions effects.

**Table 7.** Pairwise sex (male *vs* female) comparisons.

|  | Std. Error | Sig. | 95% Confidence Interval for Difference |  |
| --- | --- | --- | --- | --- |
| **Right I-IV** | 73.16 | **0.031** | 15.2 | 304.69 |
| **Right V** | 78.61 | **0.011** | 46.59 | 357.63 |
| **Right VI** | 190.8 | 0.345 | -558.37 | 196.54 |
| **Right Crus I** | 322.92 | 0.544 | -835.4 | 442.24 |
| **Right Crus II** | 256.19 | 0.085 | -62.89 | 950.72 |

Note: Post-hoc Bonferroni. Significant results are in bold.

Considering the comparisons between age groups, we observed that OA individuals exhibited significantly smaller volumes compared to the EMA group, in the right subregions I-IV, V, VI, and Crus I, indicating smaller volume with age (**Figure 8)**. Interestingly, we also observed that the LMA group had smaller volumes compared to the EMA group in the right subregions I-IV and V. No differences were observed in right Crus II. For more statistical details, see **Table 8**.

**Figure 8.**
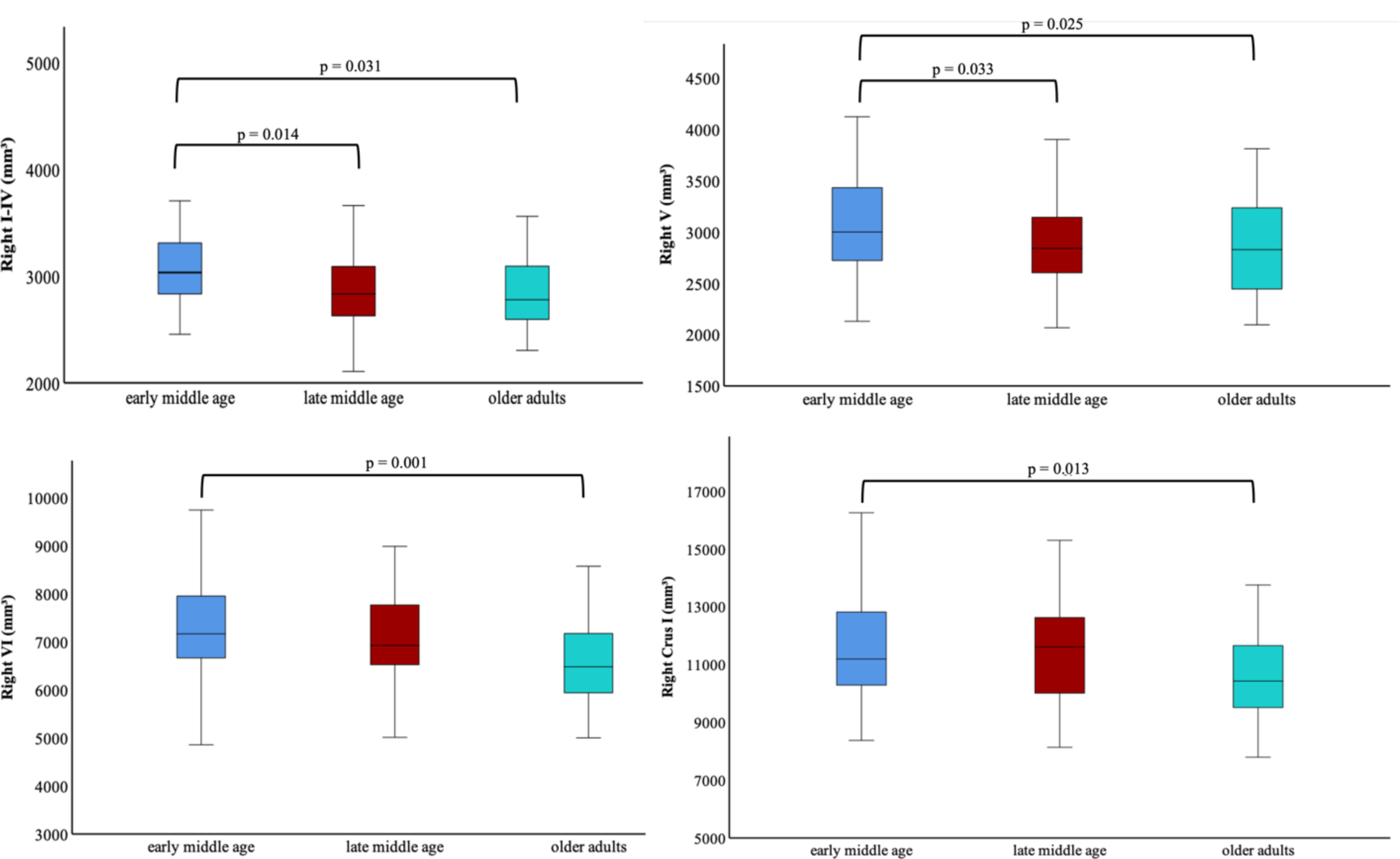
Significant age-related differences in lobular volume. The mean standard error (SE) for each group is: early middle age SE = 9.69, late middle age SE = 9.21, and older adults SE = 11.07.

**Table 8.** Pairwise age group comparisons regarding cerebellar subregions.

|  |  | Std. Error | Sig. | 95% Confidence Interval for Difference |  |
| --- | --- | --- | --- | --- | --- |
| **Right I-IV** | *LMA vs EMA* | 69.27 | **0.014** | 32.02 | 368.01 |
|  | *OA vs EMA* | 76.06 | **0.031** | 13.18 | 382.09 |
| **Right V** | *LMA vs EMA* | 74.43 | **0.033** | 11.37 | 372.38 |
|  | *OA vs EMA* | 81.72 | **0.025** | 20.73 | 417.1 |
| **Right VI** | *LMA vs EMA* | 180.65 | 0.397 | -164.63 | 711.56 |
|  | *OA vs EMA* | 198.34 | **0.001** | 256.21 | 1218.23 |
| **Right Crus I** | *LMA vs EMA* | 305.74 | 1.000 | -514.73 | 968.18 |
|  | *OA vs EMA* | 335.68 | **0.013** | 159.96 | 1788.12 |
| **Right Crus II** | *LMA vs EMA* | 242.56 | 0.401 | -222.32 | 954.14 |
|  | *OA vs EMA* | 266.31 | 0.105 | -78.9 | 1212.79 |

Note: Post-hoc Bonferroni. EMA: early middle age; LMA: late middle age; OA: older adults. Significant results are in bold.

Regarding the analysis of hormonal levels, we observed that sex and age groups are both significant while, again there was no significant sex by age group interaction (see **Table 9** for full statistical details). Males and females show significant differences in the levels of P (p=0.02) and T (p=0.001), (**Figure 9A)**. Females have higher levels of P, while males have higher levels of T.

**Figure 9.**
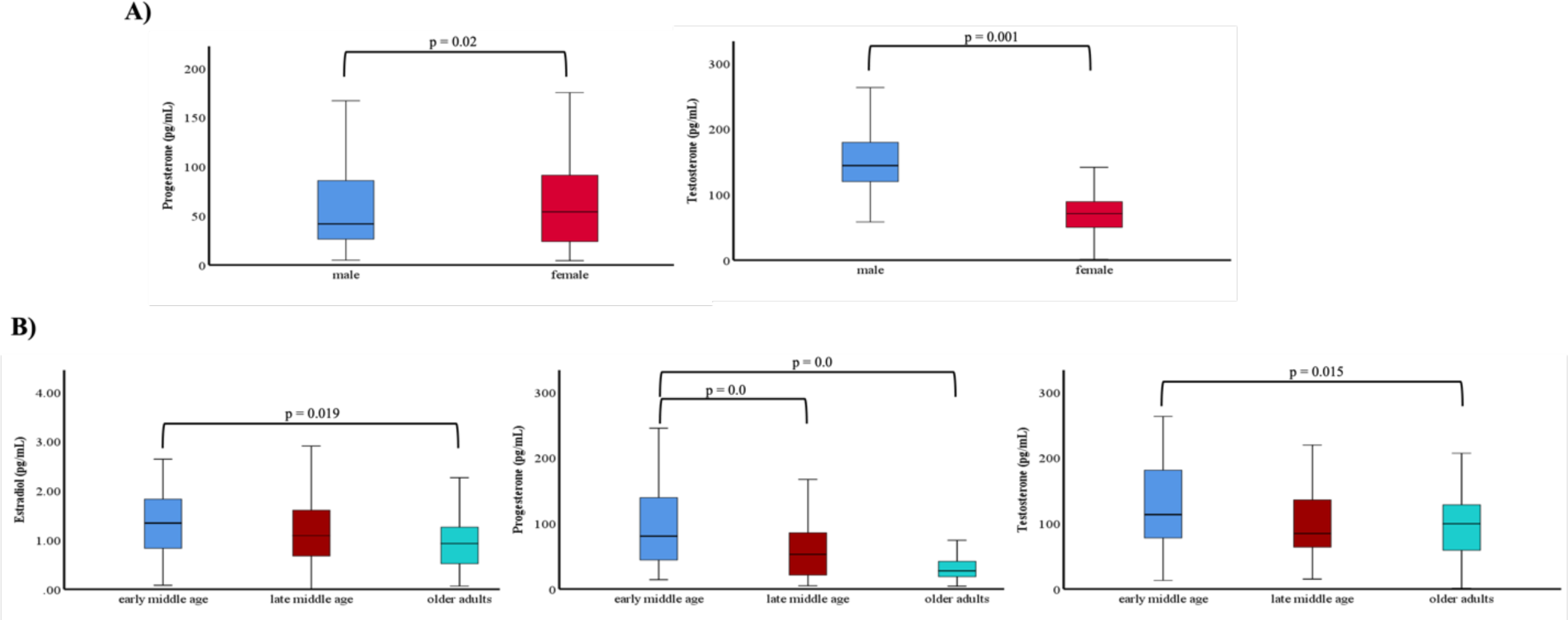
Hormonal levels analysis: A) Comparison by Sex: The standard error (SE) for progesterone is 0.74 for males and 0.72 for females, while the SE for testosterone is 0.57 for males and 0.56 for females. B) Comparison by Age. Early middle age (EMA), late middle age (LMA), and older adults (OA) estradiol SE = 0.01, 0.009, and 0.011, respectively; progesterone SE = 0.87, 0.8, and 1.0, respectively; and testosterone SE = 0.68, 0.62, and 0.77, respectively.

**Table 9.**
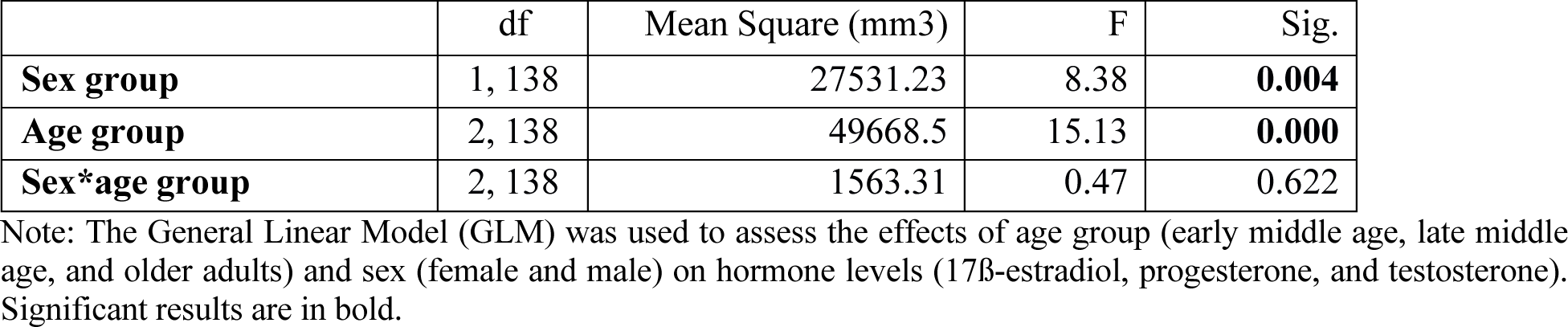
Tests of between-subjects hormone effects.

Following up on the age effect, we observed differences in E levels, with LMA females presenting lower levels compared to EMA females. For progesterone P levels, both LMA and OA females had lower levels compared to EMA females. In terms of T levels, LMA and OA males had significantly lower levels compared to EMA males. For more statistical details, see **Table 10**, and the results are illustrated in **Figure 9B**.

**Table 10.** Pairwise age group comparisons regarding hormone levels.

|  |  | Std. Error | Sig. | 95% Confidence Interval for Difference |  |
| --- | --- | --- | --- | --- | --- |
| **17β-estradiol** | *LMA vs EMA* | 0.157 | 1.000 | -0.243 | 0.519 |
|  | *OA vs EMA* | 0.176 | **0.019** | 0.060 | 0.913 |
| **Progesterone** | *LMA vs EMA* | 13.976 | **0.000** | 24.934 | 92.772 |
|  | *OA vs EMA* | 15.646 | **0.000** | 47.329 | 123.272 |
| **Testosterone** | *LMA vs EMA* | 10.860 | 0.096 | -2.788 | 49.922 |
|  | *OA vs EMA* | 12.157 | **0.015** | 5.171 | 64.179 |

Note: Post-hoc Bonferroni. EMA: early middle age; LMA: late middle age; OA: older adults. Significant results are in bold.

To better understand the relationships between hormone levels and lobular volume, we conducted linear regression analyses. The regression results indicated that T levels were significantly associated with volume in the Right I-IV and Right Crus II (**Figure 10**). These results indicate that higher T levels are significantly associated with larger volumes in the Right I-IV and Right Crus II regions. The models explain approximately 14% of the variance in the volumes of these regions, with the adjusted R-squared values accounting for the number of predictors. The positive B coefficients indicate direct relationships, and the significant t-values and p-values further confirm the statistical significance of these relationships. Statistical results are detailed in **Table 10**. No significant results were found for other hormones.

**Figure 10.**
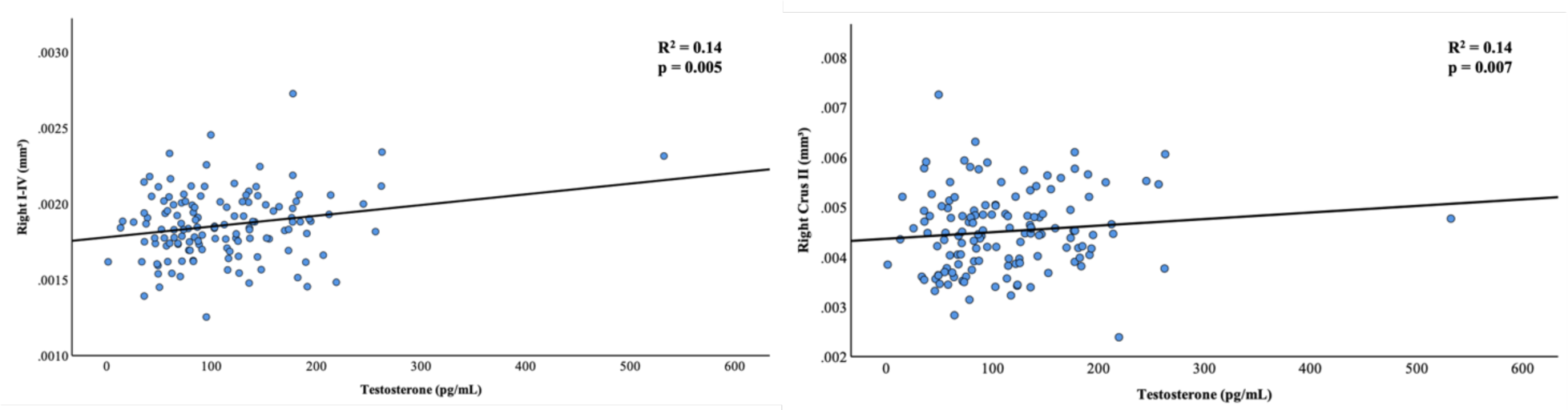
Linear regression analysis of testosterone levels predicting cerebellar subregion volumes in right I-IV and right Crus II.

**Table 10a.**
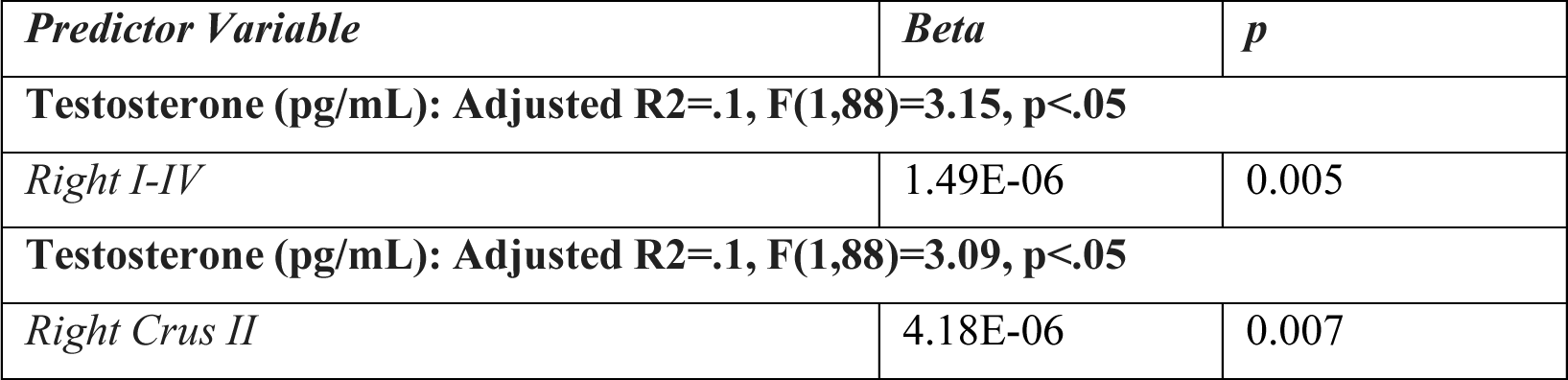
Linear regression analysis of hormone levels and lobular volume in the cerebellum subregions.

## 4. Discussion

In this study, we conducted a novel investigation into the relationship between sex, sex steroid hormone levels, regional cerebellar volume, and resting-state connectivity in middle-aged and older adults. Consistent with prior research, we observed age-related differences in hormone levels. Specifically, our results indicated higher 17β-estradiol levels in the EMA group compared to the OA group, aligning with the typical decrease in 17β-estradiol levels with age, particularly during menopause [6, 46]. Additionally, we noted differences in progesterone and testosterone levels between the EMA group and both the LMA and OA groups. Furthermore, we identified sex differences in cerebellar lobular volume and connectivity, as well as significant relationships between sex steroid hormones and these cerebellar metrics. The implications of these findings are discussed in more detail below, within the context of the broader literature.

Regarding our findings on differences in brain connectivity between males and females, several notable patterns emerged. Females exhibited less connectivity between lobules I-IV and the cuneus compared to males as they aged, suggesting a potential divergence in neural communication patterns between the sexes with increasing age. This reduced connectivity in females may influence sensory processing and integration, which are key functions of these cortical regions [27, 47]. Conversely, females showed greater connectivity between Crus I, Crus II, and the precuneus compared to males as they aged, aligning with previous studies [48]. Notably, a parallel study by our group (Tracey’s paper), revealed associations between higher Crus I and superior parietal (anatomically just posterior to the precuneus) connectivity to better performance on a composite measure of cognitive tasks broadly assessing episodic memory, executive function, and motor learning; however, these associations were found across both female and male participants. The higher connectivity seen here paired with our group’s cognitive correlational [32] findings suggests that females may maintain connectivity in these regions, which are involved in higher-order cognitive processes such as visuospatial imagery, episodic memory retrieval, and self-referential thinking [32, 49, 50]. Alternatively, this could be an attempt at compensation for other age-related processes. Additionally, we found that females have lower connectivity between lobule V and the calcarine cortex than males with increasing age. This could indicate sex differences in the visual processing pathways, potentially affecting visual perception and processing speed as females age [11, 51, 52].

It is important to highlight that females may face greater consequences with age in terms of organization and efficiency of functional networks. This could contribute to the disproportionate impact of normative behavioral declines and age-related illnesses on women’s aging compared to males. However, as discussed in the work of Ballard et al. (2022) [48], the differences observed in cerebellar connectivity between reproductive and postmenopausal women may be attributed to hormonal fluctuations. Indeed, dense sampling studies of the menstrual cycle have highlighted associations between hormonal fluctuations and functional network organization in the young female brain [23, 53]. While the menopausal differences implicated here occur on a different time scale, there may be impacts of these hormonal fluctuations.

Relatedly, our hormonal analyses found that higher 17β-estradiol levels were linked to higher connectivity between Crus I and the superior frontal gyrus, and between Crus II and the cuneus, suggesting enhanced interactions with these regions with higher levels of hormone. 17β-estradiol has been previously implicated in modulating brain connectivity [46, 54] and enhancing cognitive performance [49], particularly in tasks requiring memory and executive function. In contrast, higher testosterone levels showed a dual role in brain connectivity. Higher testosterone was linked to greater connectivity between lobules I-IV and both the angular and lingual gyri. The angular gyrus is known for its role in complex language functions, mathematical processing, and spatial cognition [55]. The lingual gyrus is involved in visual processing, particularly in recognizing words and faces [56]. The observed connectivity in these regions suggests that testosterone may support and enhance these specific cognitive processes [57–59].

However, testosterone was also associated with lower connectivity between Crus I and the middle temporal gyrus, the superior frontal gyrus, and lobule V. The middle temporal gyrus is critical for semantic memory and language processing [60]. The superior frontal gyrus is involved in higher cognitive functions, including working memory and attention [61], while lobule V plays a role in motor control and coordination [62]. Lower connectivity in these regions might indicate that high testosterone levels influence these cognitive and perceptual processes, potentially leading to variations in cognitive performance and perceptual abilities [58]. Previous studies have shown that testosterone influences brain structure and function, impacting cognitive performance and behavior [25, 63, 64]. For example, one study found that testosterone administration in aging men improved spatial memory and executive function [25]. Another study reported that endogenous testosterone levels were associated with variations in brain volume and connectivity in regions involved in language and visuospatial abilities [65].

Lastly, our analysis revealed that higher progesterone levels were associated with reduced connectivity between Crus I and the putamen in females compared to males. Additionally, females with higher progesterone levels exhibited reduced connectivity between lobule V and the middle temporal gyrus. Progesterone, known for its neuroprotective effects and modulation of neurotransmitter systems crucial for motor control and cognitive processes, underscores the need for further research to understand its role in modulating neural circuits and its implications for neurological disorders and cognitive function [66]. Therefore, the sex-specific effects of progesterone on brain connectivity highlight the need for further research to better elucidate its role in modulating neural circuitry and its potential implications for neurological disorders and cognitive function [67]. Recent studies have suggested a potential role for testosterone in cognitive function and brain health in women as well [58, 68, 69]. It is conceivable that higher testosterone levels may influence brain connectivity patterns, highlighting the need to understand its interactions with other sex hormones like progesterone and estradiol. This understanding could provide valuable insights into the complex interplay of hormones in the aging brain, particularly in females. This may be particularly important in post-menopausal females when there are very low levels of 17β-estradiol and progesterone, and testosterone effects may become more prominent. That is, shifts in the hormonal balance may result in more robust associations between testosterone and both brain and behavioral metrics. Further research could clarify whether testosterone plays a protective or detrimental role in brain aging in females, and certainly, more work in this area is needed.

Significant differences in cerebellar subregion volume were observed with age and between sexes, particularly in the right subregions I-IV and V, where males had larger volumes compared to females. EMA individuals had larger volumes compared to LMA and OA in the same regions, as well as in the right Crus I and VI regions. Although there are several studies showing cerebellar volume differences as individuals age [70–72], studies focusing on lobular volume have been more limited. Previous studies from our group have addressed this issue and have also shown differences in the right Crus I, V, and VI between young and old individuals, with young people also having larger volumes [15, 24]. Although we did not find significant interactions between sex and age, we observed that EMA has higher estrogen levels and larger cerebellar subregion volumes, and we speculate that this could be related to the neuroprotective effects of estrogen [15] and testosterone (discussed in more detail below). Estrogen has been shown to promote neuronal growth and synaptic plasticity [47]. However, as individuals age, particularly in females the decline in estrogen levels, particularly post-menopause, may lead to accelerated volume reduction [50]. As hormone levels fluctuate with age – decreasing in both sexes, but at different rates and patterns – the impact on brain structure and connectivity can become more pronounced [4, 24, 48].

In relation to our regression analysis findings, we observed positive relationships between testosterone levels and lobular cerebellar volume. These associations suggest that testosterone may play a crucial role in maintaining cerebellar structure, which could have implications for motor and cognitive functions mediated by these regions. Furthermore, testosterone’s ability to predict cerebellar volume highlights the potential for hormone levels to serve as biomarkers for brain health and aging [25, 64, 68]. The significant relationship between testosterone and cerebellar volume may also reflect the broader neuroprotective effects of testosterone [69], as suggested by previous studies linking testosterone to brain volume and connectivity in regions involved in cognitive and motor functions. Further, as noted above, relationships with testosterone may be more notable, particularly in older females as there are changes in the broader hormonal environment and balance, though further work is certainly needed. Notably, there were no associations with 17β-estradiol and progesterone. 17β-estradiol in particular is at very low levels in OA females, and progesterone is also significantly lower. As such, there was much more limited variability in these measures in our sample (and males were included, where again levels are lower), perhaps making these associations more difficult to assess, if present at all. Future follow-up work looking at changes in these hormones over time in younger females concerning lobular volume stands to be particularly informative.

While our study’s limitations include the use of cross-sectional data, which prevents us from making causal inferences or understanding individual changes over time, it underscores the significant role of sex hormones in modulating brain connectivity and structure across different age groups. The observed differences in connectivity and brain volume between males and females, as well as among various age groups, emphasize the importance of considering sex and hormonal status in neuroimaging studies. It is also crucial to recognize that hormone levels can vary significantly both within a single menstrual cycle and between different females [73]. Additionally, our sample includes both regularly cycling reproductive females, those transitioning into menopause, and post-menopausal females. Longitudinal studies of sex hormone levels in the same participants over time could clarify individual variations in hormonal change. Our findings enhance our understanding of how hormonal differences associated with aging influence brain functional networks and structure, providing insights that could inform future research and potential therapeutic strategies for age-related cognitive decline and neurological disorders.

## Acknowledgment

This work was supported by R01AG065010 to J.A.B. This work was further supported by the Texas Virtual Data Library (ViDaL), a high-performance cluster, funded by the Texas A&M University Research Development Fund. In this cluster, the imaging analyses for the current work were carried out using the resources provided by the Texas A&M High-Performance Research Computing organization.

